# High-Yield Induced Ovulation in Adult Fat-Tailed Dunnarts by PMSG Treatment Combined with Estrus Synchronization

**DOI:** 10.1101/2025.08.19.671172

**Authors:** Jun Liu, Namdori Mtango, Emily L. Scicluna, Sara Ord, Andrew J. Pask

## Abstract

The fat-tailed dunnart, *Sminthopsis crassicaudata*, is an emerging laboratory-based marsupial model for research on comparative biology, reproduction and conservation. In females, the reproductive cycle lasts 31 days and approximately 10 oocytes are ovulated per cycle. Assisted reproductive technologies (ART) play a crucial role in marsupial conservation, but developing protocols to harvest large numbers of oocytes remains a key challenge. Specifically, producing sufficient mature metaphase II (MII) oocytes in dunnarts continues to be difficult. Ovarian follicle stimulation is common practice to achieve superovulation in many species and typically requires treatment of prepubertal female animals to avoid the impacts of endogenous hormone cycling. Alternatively, adult females can be stimulated during the intermediate or follicular phase. In this study, we aimed to develop a high-yield induced-ovulation protocol to collect higher number of MII oocytes from adult, cycling, female dunnarts. We first synchronized the female dunnart reproductive cycles using luteinizing hormone-releasing hormone (LHRH). The reproductive cycles were monitored by examining cytology of vaginal lavage samples. After administering four LHRH injections given at three-day intervals, 88.9% (n=36) of the adult female dunnarts responded to the treatment, with their estrous cycles synchronized at the diestrous stage. We then induced ovarian follicle development through two PMSG injections over 6 days, followed by hCG administration to trigger ovulation. By combining estrous cycle synchronization with PMSG stimulation, we achieved 77.8% (n=36) ovulation success and obtained an average of 20.1±9.1 (n=28) MII oocytes per adult dunnart. These data demonstrated that estrous cycle synchronization followed by the PMSG-hCG treatment yields consistent, highly efficient induced-ovulation in adult dunnarts. This approach of combining estrus synchronization and follicle stimulation to produce sufficient MII oocytes for ART purpose could be applied to other valuable marsupial species to support conservation efforts.

**Summary Sentence:** Superovulation and robust production of mature MII oocytes can be induced by 10-day estrus synchronization using LHRH followed by 6-day PMSG stimulation of follicle growth in adult fat-tailed dunnarts, a marsupial species.

**Graphical Abstract:** 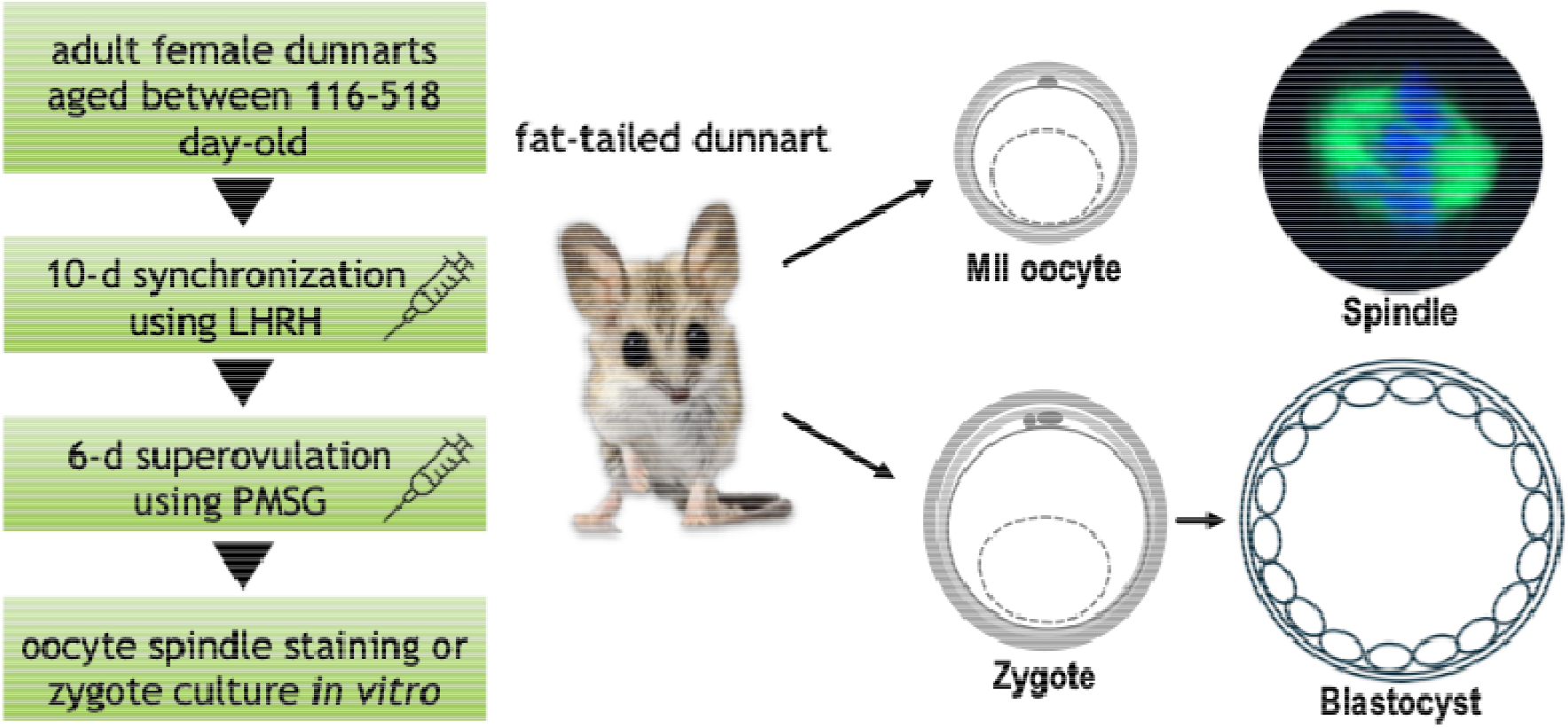

## Introduction

Harvesting sufficient mature metaphase II (MII) oocytes is critical for the use of assisted reproductive technologies (ART), including in vitro fertilization (IVF), intracytoplasmic sperm injection (ICSI), and somatic cell nuclear transfer (SCNT) or cloning, in the fat-tailed dunnart (*Sminthopsis crassicaudata;* hereafter referred to as the dunnart) reproduction and conservation studies. Recently, we developed a robust superovulation protocol to generate MII oocytes in prepubertal dunnarts [1]. This protocol involves three injections of pregnant mare serum gonadotropin (PMSG) over 6 days to stimulate ovarian follicle development, followed by an injection of human chorionic gonadotropin (hCG) to trigger ovulation. While prepubertal induced ovulation is standard for many model animal species, this may not be practical when working with wildlife or animals with complex husbandry. However, when we applied the prepubertal regimen to adult female dunnarts, both the ovulation rates and oocyte yields varied substantially between animals.

Superovulation in prepubertal mice is preferentially used for oocyte and embryo collections due to the higher yield and better quality of the mature oocytes and zygotes [2]. Studies in B6 and DD mice have shown that the estrous cycle stage significantly affects superovulation efficiency in adult animals [2, 3]. In adults, gonadotropin-releasing hormone (GnRH) production is released in a characteristic pulsatile pattern. This initiates the functioning of the hypothalamic-anterior pituitary-gonadal (HPG) axis [4]. When administering PMSG for stimulating follicle development in adults, the timing rarely aligns with their natural estrous cycle. This misalignment typically creates conflicting signals between endogenous hormones and injected exogenous hormones. Previous gonadotropin-induced estrus and ovulation in fat-tailed and stripe-faced dunnarts produced limited and inconsistent outcomes [5-8].

Alternatively, when stimulated during the intermediate or follicular phase, the adult animals produced superior results [8, 9]. GnRH analogs have been used for decades to synchronize estrus in farm animals for the improvement of reproductive efficiency [10, 11]. In laboratory rats, luteinizing hormone releasing hormone agonist (LHRHa), a synthetic analogue, could synchronize the estrous cycle [12]. In fat-tailed dunnart experiments, Lucrin Depot (a 1-month microsphere GnRH agonist preparation) suppressed reproductive activity for 4-8 weeks, with activity returning at 8-12 weeks [13, 14]. This approach showed potential as an assisted breeding tool, as pouch young were born to two females following the treatment [13, 14]. However, it remains unclear whether this approach need to be combined with ovarian superovulation treatment for effective sufficient oocyte collection in the fat-tailed dunnarts.

In this study, we developed an effective protocol for harvesting mature MII oocytes from adult female dunnarts. We first established a method for estrous cycle synchronization using LHRH and then evaluated whether the synchronized females responded well to the 6-day PMSG-hCG superovulation protocol [1]. To assess the functional normality of oocytes produced through the cycle synchronization and superovulation combined approach, we conducted *in vivo* fertilization and *in vitro* zygote culture experiments. This high-yielding protocol combining estrus synchronization and superovulation in adult females could enhance the efficacy of ART implementation in dunnarts. Furthermore, this approach could potentially support conservation efforts for other marsupial species, particularly those with reproductive challenges.

## Materials and Methods

### Animals

Experiments in this study were performed using adult fat-tailed dunnarts (116 - 518 days of age) housed at the BioSciences 4 animal facility, University of Melbourne. Animals were maintained at 25°C with 50% humidity under controlled lighting conditions (16 h light from 5:00 am to 9:00 pm and 8 h dark from 9:00 pm to 5:00 am) [15]. The Animal Research Ethical Committee of the University of Melbourne approved all procedures in this study (AEC26863 and AEC26864).

### Vaginal lavage cell staining

Female dunnart vaginal lavage samples were collected and stained as previously described [1]. In brief, female dunnart vaginal lavage samples were collected in 20 μL of phosphate buffered saline (PBS) from the vaginal opening, transferred to slides, air-dyed and stained with 0.1% crystal violet (Sigma). The stained lavage samples were examined under a light microscopy (Motic) to assess cell types and quantities of nucleated epithelial cells (NEC), cornified epithelial cells (CEC) and polymorphonuclear leukocytes [16].

### Estrus synchronization in dunnarts

Female dunnarts received subcutaneous injections of 20 μg (0.1 mL) of LHRH (D-Ala^6^-LH-RH acetate salt hydrate, Sigma) per animal, once every 3 days for 10 days (Figure 1). Vaginal lavage samples were collected the day after each of the LHRH injections to monitor estrous cycle status through lavage cytology [17, 18]. The estrous cycle stages were classified according to the following criteria based on the cell types and quantity in the lavage samples: proestrus was characterized by the presence of nucleated epithelial cells with few cornified epithelial cells and/or leukocytes; estrus by predominance of polygonal cornified epithelial cells; metestrus by an influx of leukocytes with few cornified epithelial cells and/or nucleated epithelial cells; and diestrus by decreased numbers of all three cell types.

**Figure 1.**
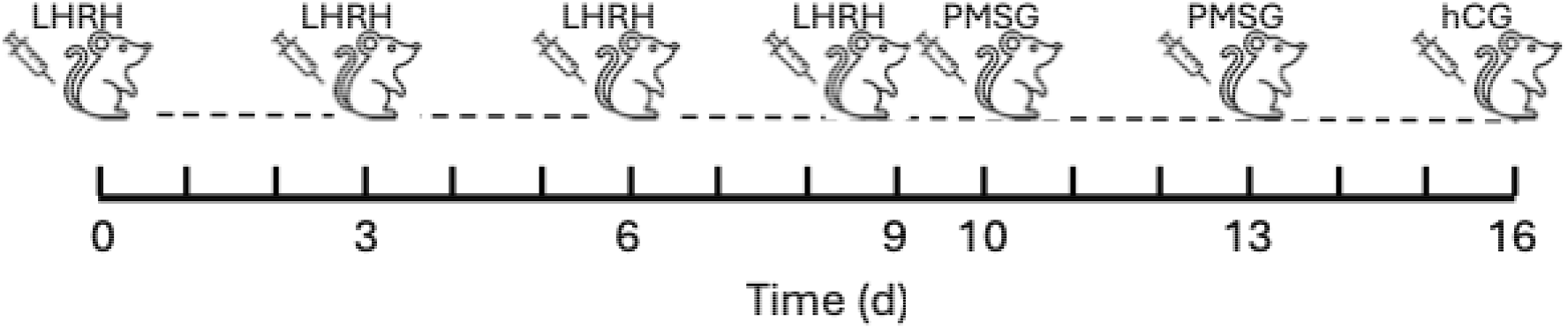
A timeline scheme for synchronization and superovulation. Female dunnarts were injected 4 times with 20 µg LHRH per animal and primed with 1 IU PMSG twice, on days 10 and 13, followed by hCG injection on day 16 to trigger ovulation.

### Superovulation and oocyte collection

We applied a 6-day dunnart superovulation protocol as we previously developed in prepubertal dunnarts [1] with modifications in this study. Female dunnarts that underwent estrous cycle synchronization received subcutaneous injections of 1 IU PMSG (ProSpec, Ness Ziona, Israel) on day 10 and day 13, followed by an intraperitoneal injection of 1 IU hCG (ProSpec) on day 16 (Figure 1). We collected mature MII oocytes from oviducts and uteri 10-14 hours after hCG administration. The oviducts and uteri were dissected free and transferred to CO_2_ independent medium supplemented with 10% fetal bovine serum (FBS), 10 IU/ml penicillin, and 100 μg/ml streptomycin (hereafter referred to as handling medium; all products from ThermoFisher). Oocytes were flushed out from the uteri and oviducts using a 30G needle with handling medium, and the number of ovulated oocytes was counted.

### Spindle fluorescence staining

Oocytes were fixed in 4% paraformaldehyde (Sigma) in PBS (pH 7.4) for 40 min at room temperature and permeabilized in 0.5% Triton X-100 (Sigma) for 20 min at room temperature. The oocytes were then blocked in PBS supplemented with 1% BSA, 0.1% Tween 20 (Sigma) and 0.01% Triton X-100 (PBST) for 1 hour before being incubated with anti-α-tubulin-FITC antibody (Sigma, 1:200 dilution) at 4 °C overnight. After washing in PBST, the oocytes were counterstained with 10 µg/ml Hoechst 33342 (Sigma) for 10 min. Finally, the oocytes were transferred to wells on a μ-Slide (Ibidi) and observed under a laser scanning confocal microscope (Nikon).

### Zygote Retrieval and in vitro culture

Female dunnarts undergoing synchronization and superovulation were paired with stud male dunnarts in a 1:1 ratio after the second PMSG injection. We collected zygotes from oviducts and uteri 5-6 hours after hCG administration. After washing 3 times in handling medium, we incubated 4-5 zygotes in 30 μL drops of dKSOM1 medium for 3 days and then transferred them to dKSOM2 medium for blastocyst formation. The compositions of the dKSOM1 and dKSOM2 medium were described elsewhere [1, 19].

### Data analysis

Percentages were calculated from more than three independent replicate experiments. The percentages were compared between groups using chi-square analysis with data pooled from all experiments in the same group. Oocyte numbers were presented as mean ± SD (standard derivation) and compared using Student’s *t*-test. Differences were considered significant when *P* ≤ 0.05.

## Results

### Vaginal lavage cytology and estrous state

We first examined cell types and their relative numbers in vaginal lavage samples to determine correlations with the reproductive (or estrous) cycle stage. Using the crystal violet staining method, three primary cell types were detected in vaginal lavage samples: nucleated epithelial cells (NEC, Figure 2A), enucleated cornified epithelia cells (CEC, Figure 2B) and polymorphonuclear leukocytes (Figure 2C). Leukocytes appeared small in size with darkly stained polymorphic nuclei. CECs were distinguished from NECs by their large irregular or polygonal shape and the absence of a nucleus. During proestrus, samples contained NECs often with a combination of the other two cell types (Figure 2D). Estrus was characterized a predominance of polygonal CECs, with a significantly elevated number of CECs often appearing in clusters (Figure 2E). Metestrus showed an influx of leukocytes with few CECs and/or NECs (Figure 2F). During diestrus, the numbers of all three cell types were significantly decreased compared to the other stages (Figure 2G).

**Figure 2.**
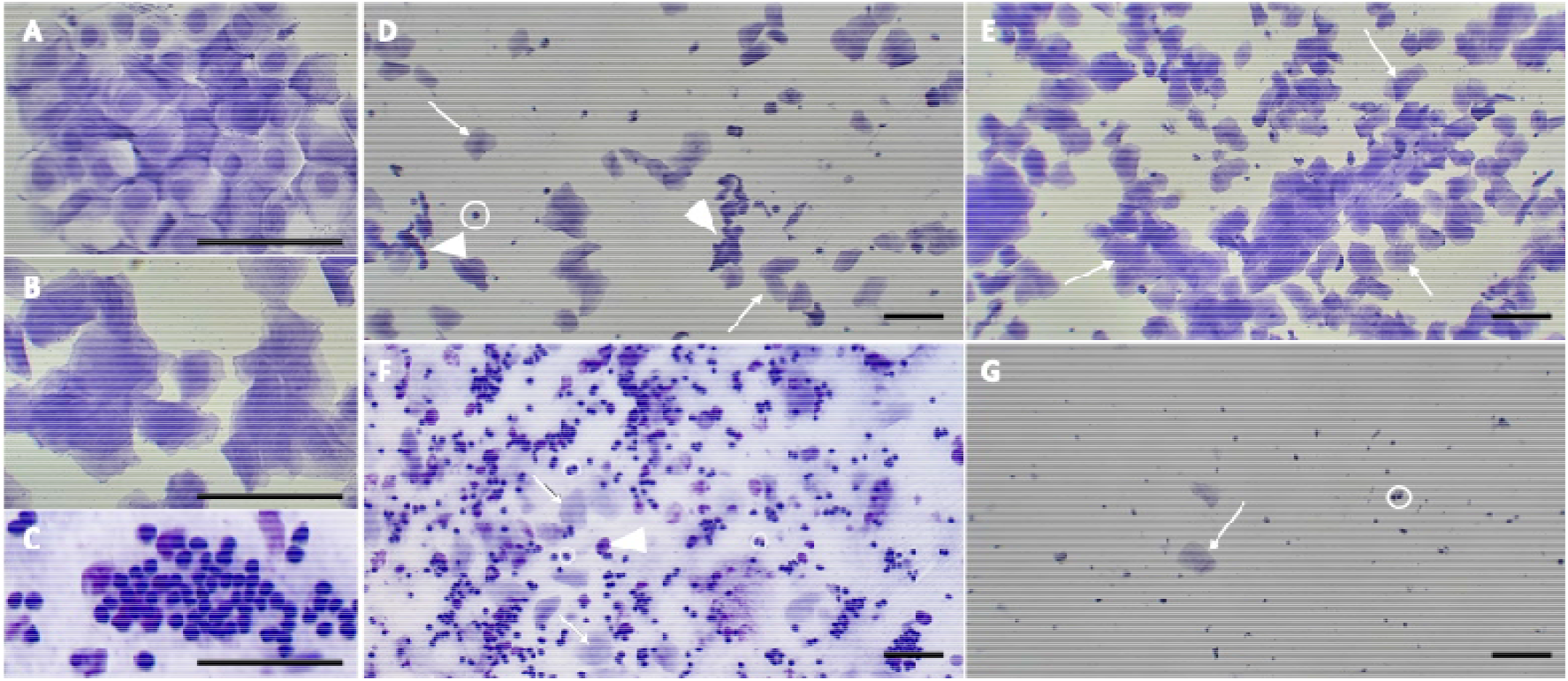
Representative images of crystal violet-stained vaginal lavage cytology. Three primary cell types observed in vaginal lavage: A) nucleated epithelial cells (NEC), B) cornified epithelial cells (CEC), and C) leukocytes. The four estrous stages include: D) Proestrus, showing a mixture of NEC (arrowheads), leukocytes (circle), and CEC (arrows); E) Estrus, characterized by predominant CEC (arrows); F) Metestrus, displaying an influx of leukocytes with some CEC (arrow) and few NEC (arrowhead); and G) Diestrus, showing markedly reduced numbers of leukocytes (circle) and CEC (arrow). Bar=100 µm.

### Estrous cycle synchronization

We then examined whether the estrous cycles of adult female dunnarts at various stages of their reproductive cycle could be synchronized through sequential LHRH injections on Days 0, 3, 6 and 9. By Day 10, 88.9% (n=36) of the dunnarts had entered diestrus (Table 1). This indicates that a 10-day continuous LHRH treatment protocol (20 μg/animal/injection) administered via four subcutaneous injections was effective for estrus synchronization.

**Table 1.** Reproductive cycle stages of adult dunnarts (n=36) following LHRH injections on Days 0, 3, 6 and 9.

| Estrous stage | No. of females (%) |  |  |  |
| --- | --- | --- | --- | --- |
|  | Day 1 | Day 4 | Day 7 | Day 10 |
| Proestrus | 16 (44) | 16 (44) | 9 (25) | 1 (3) |
| Estrus | 2 (6) | 2 (6) | 0 (0) | 1 (3) |
| Metestrus | 3 (8) | 1 (3) | 2 (6) | 2 (6) |
| Diestrus | 15 (42) | 17 (47) | 25 (69) | 32 (89) |

### Oocyte yield and maturation

For generation and harvesting mature MII oocytes in adult dunnarts, we examined the combined effects of estrous cycle synchronization with LHRH treatment and superovulation with PMSG treatment. Following the combines treatments, we examined the number of females that ovulated, and oocytes retrieved. Among 36 adult female dunnarts (aged 116-518 days) subjected to the combined synchronization and superovulation treatments, 28 (77.8%) successfully ovulated with oocytes collected post-hCG administration. This rate was significantly higher than the control group (*P* = 0.037; Figure 3A). In the control group, only three (37.5%) of eight adult females that were superovulated with the 6-day PMSG treatment without prior synchronization ovulated. The average yield of MII oocytes in the synchronization group was 20.1±9.1 (n=28) per adult female, which was significantly higher than that in the control group (5.7±2.3, n=3; *P* = 0.012). The normal MII oocytes displayed the first polar body in the thin perivitelline space and a characteristic deutoplasm vesicle in the ooplasm and were surrounded by zona pellucida and a layer of mucoid coat (Figure 3B). Oocytes immunostained with anti-⍰-tubulin-FITC antibody and counterstained with Hoechst 33342 revealed a barrel-like spindle apparatus with well-aligned chromosomes on the metaphase (Figure 3C).

**Figure 3.**
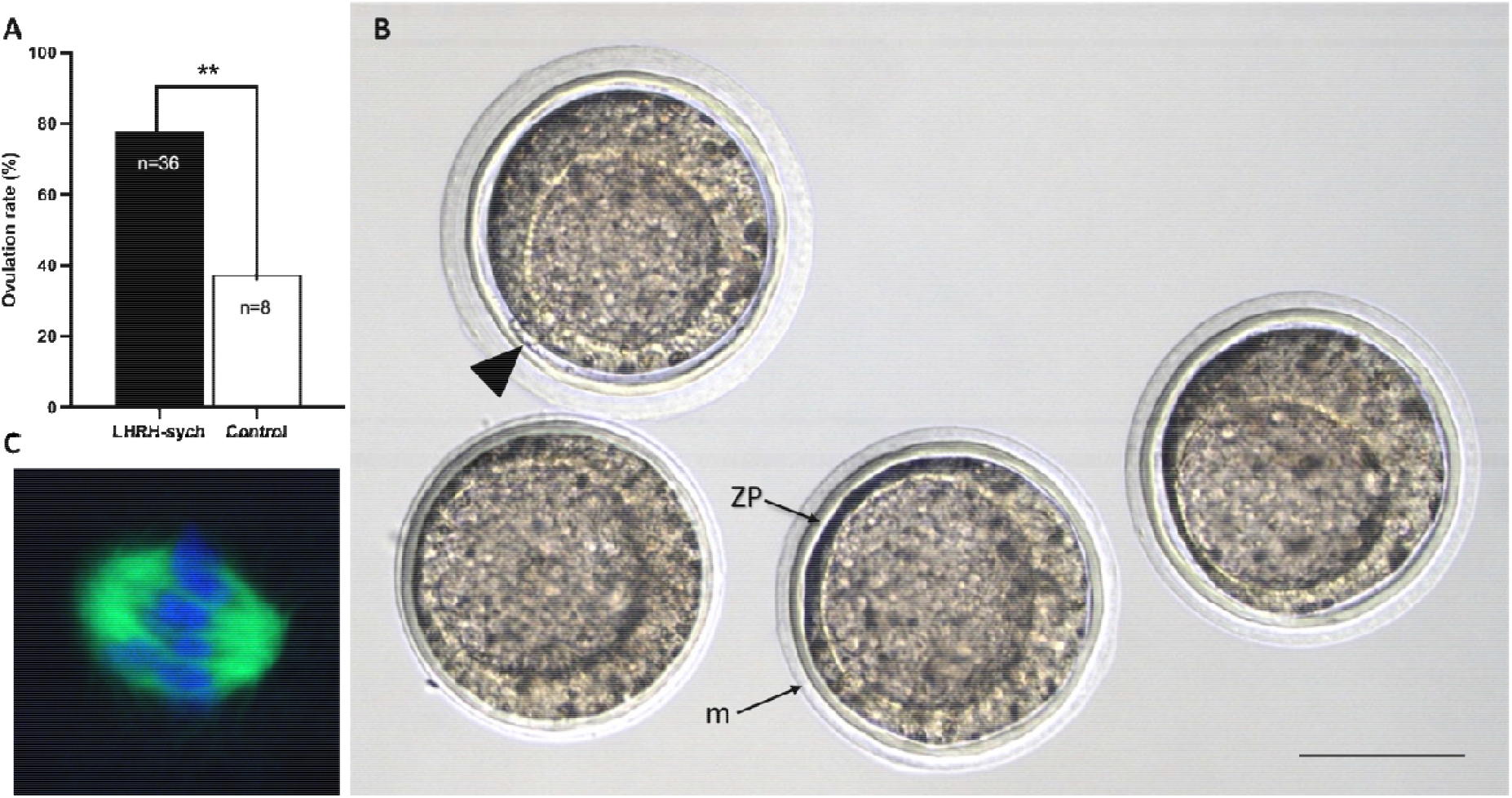
Dunnart ovulation rates and oocyte characterization. A) Ovulation rate in the LHRH-treated females was higher than that in control group (P = 0.037). B) MII oocytes with the first polar body (arrowhead) in the thin perivitelline space and the characteristic deutoplasm vesicle in ooplasm, surrounded by zona pellucida (ZP) and mucoid coat (m). C) A representative image of spindle morphology and chromosome alignment in an oocyte at the MII stage. Bar=100 µm.

### *In vivo* fertilization and zygote culture

To assess the developmental capacity of the MII oocytes produced from adult female dunnarts after combined synchronization and superovulation treatments, we performed *in vivo* fertilization by mating the treated females with stud male dunnarts. Zygotes were collected and cultured in dKSOM1/dKSOM2 sequential medium to evaluate early embryo development competence. The results are summarized in Table 2. Two of 4 treated dunnarts were successfully mated, and a total of 49 zygotes retrieved (Figure 4A). A remarkable feature of dunnart zygotes is that multiple sperms were trapped in and inside the mucoid layer. After 48 hours of culture in dKSOM1 medium, 87.8% (n=49) zygotes developed to 4-cell embryos. Interestingly, these embryos paused at the 4-/5-cell stage by the next day (Figure 4B). Embryos were then transferred to dKSOM2 medium and 76.7% (n=43) of 4-cell embryos developed to 8-/16-cell embryos in 24 hours in dKSOM2 medium (Figure 4C). At approximately 120 h post-hCG administration, 51% (n=49) zygote embryos had developed to the blastocyst stage (Figure 4D).

**Table 2.** Developmental capacity of dunnart zygotes fertilized in vivo.

| Animal # | Successful mating | No. of oocytes | No. of zygotes | No. (%) of embryos developed into* |  |
| --- | --- | --- | --- | --- | --- |
|  |  |  |  | 4-cell | Blastocyst |
| 1 | No | 22 | 0 | n.a. | n.a. |
| 2 | No | 24 | 0 | n.a. | n.a. |
| 3 | Yes | 0 | 20 | 19 (95) | 12 (60) |
| 4 | Yes | 0 | 29 | 24 (83) | 13 (45) |
| Total |  | 46 | 49 | 43 (88) | 25 (51) |

\*: percentages calculated based on number of zygotes.

**Figure 4.**
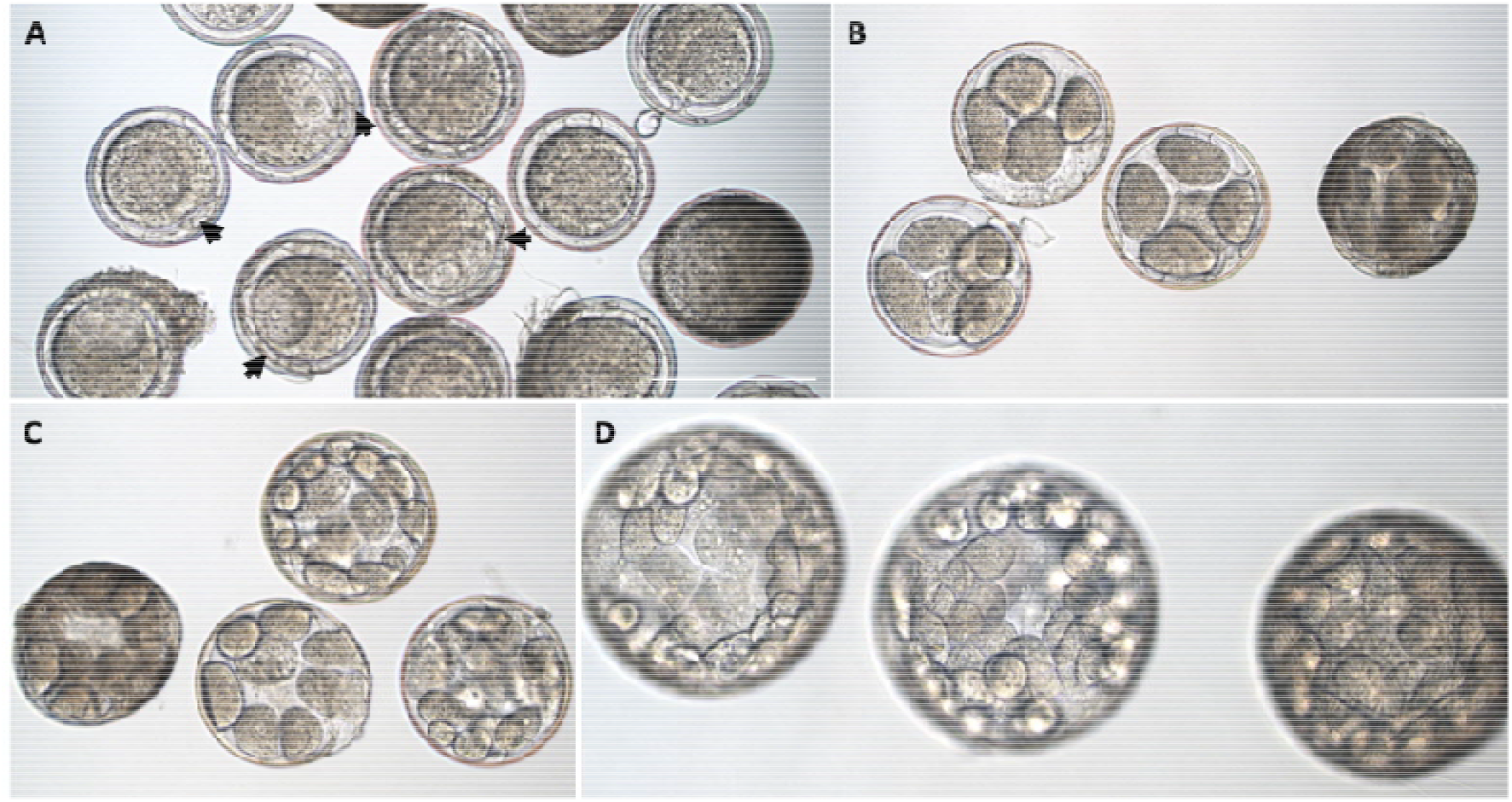
Dunnart zygote culture in vitro. (A) Zygotes collected 5-6 h post-hCG with multiple sperms in and inside mucoid layer (arrows), which (B) developed into 4-cell stage embryos at 48-72 h post-hCG, (C) progressed to approximately 8-/16-cell stage embryos at 96 h post-hCG, and (D) formed blastocyst embryos at about 120 h post-hCG when cultured in sequential dKSOM1/dKSOM2medium. Bar=250 µm.

## Discussion

The fat-tailed dunnart, *Sminthopsis crassicaudata*, is an emerging laboratory-based marsupial model for research on comparative biology, reproduction and conservation. This study was undertaken to establish high yield induced ovulation protocols for adult female fat-tailed dunnarts. In females, the reproductive cycle lasts about 31 days and approximately 14 oocytes are naturally ovulated [20, 21]. Producing sufficient mature MII oocytes remains a challenge in dunnarts, yet they are crucial for developing assisted reproductive technologies (ART) for marsupials. Ovarian follicle stimulation is commonly used to achieve superovulation across a broad range of species and is typically performed in prepubertal females to avoid the impacts of endogenous hormone levels. We confirmed that our recently developed protocol enabled us to successfully induce ovulation from prepubertal dunnarts aged 70-85 day old for the first time [1]. However, when we applied the prepubertal superovulation regimen to adult female dunnarts, both the ovulation rates and oocyte yields were inconsistent and varied substantially between individuals. The variability in reproductive outcomes reflected the different responses of each animal to the PMSG stimulation due to their varying stages in the estrous cycle. Studies in strip-faced dunnarts has shown that adult animals stimulated during the intermediate or follicular phase would produce superior results [8, 9]. To achieve this, we first synchronized the adult female dunnart reproductive cycle using luteinizing hormone-releasing hormone (LHRH). Reproductive cycle status of the females was monitored and determined by cytology of vaginal lavage samples. After four LHRH injections administered every three days, 88.9% (n=36) of the adult female dunnarts responded to the LHRH treatment, with estrous cycles synchronized at the diestrous stage of reproductive cycle. We then induced ovarian follicle development through two PMSG injections for 6 days, followed by hCG administration to trigger ovulation. By combining estrous cycle synchronization and PMSG stimulation, we achieved 77.8% (n=28) ovulation rate and obtained 20.1±9.1 (n=28) MII oocytes per adult dunnart. Thus, this demonstrated that estrous cycle synchronization followed by the PMSG-hCG treatment yielded consistent, highly efficient induced-ovulation in adult dunnarts. For conservation purposes and captive management, prepubertal animals are not always suitable or available for manipulation, therefore developing a robust superovulation protocol for adult female dunnarts has practical significance. This new methodology also increases the yield of developmentally competent oocytes and embryos per animal, further facilitating the development of ART technologies for marsupials.

For the superovulation part of the protocol, we modified the original sequence of the three intraperitoneal (IP) PMSG injections (every 2 days) [1] with two subcutaneous (SC) injections administered 3 days apart in this study. The total of 6-days PMSG priming was unchanged but the reduced injection number helps to reduce stress which could introduce endocrine noise to the study [22]. Secondly, SC absorption is deliberately slower than IP, creating a depot effect that maintains systemic gonadotropin concentrations across the longer interval [23]. Additionally, switching to SC might also remove the 10-20% “mis-injection” error and the risk of visceral puncture associated with IP delivery, thereby improving animal welfare and reducing inter-animal variation [24, 25]. Together, the two-dose SC regimen refines the protocol by lowering animal stress, minimising procedure risk and taking advantage of the SC-longer hormone coverage benefit. This improved method is now also applied in our routine experiments for oocyte and embryo generations in prepubertal dunnarts.

The effect of estrous cycle stage on superovulation efficiency has been investigated in rats and mice, which have a 4–5-day estrous cycle length, with varied results [3, 26-28]. Guinea pigs, which is an animal with complete estrous cycle and have 16-18 day cycle length, could be induced to undergo synchronized ovulation via progesterone treatment [29]. Studies in the stripe-faced dunnarts (*Sminthopsis macroura*), demonstrated superior outcomes when females were stimulated during the intermediate or follicular phase, avoiding the luteal phase period [8, 9]. The dunnart is another animal with a complete estrous cycle of about 30-31 days since its corpora lutea persists through most of the luteal phase [20]. Witt et al reported that adult dunnarts could undergo synchronized ovulation through subcutaneous injection of Lucrin Depot, a GnRH agonist [13, 14]. The GnRH secretion in a pulsatile pattern is fundamental for mammalian reproduction [30, 31]. The continuous GnRH administration in animals paradoxically inhibits gonadotropin release [32, 33]. Studies have shown that long-acting GnRH treatment causes receptor overstimulation, resulting in receptors losing their GnRH affinity, or becoming desensitized [32]. This desensitization reduces circulating FSH and LH levels. The GnRH-induced gonadotropin downregulation inhibits ovarian function, leading to impaired follicular growth and development during treatment [4, 34, 35]. In this study, we first demonstrated that the estrous cycle in dunnarts could be synchronized simply by administering LHRH for 10 days. Most dunnarts (89%) reached the diestrus stage by Day 10 after LHRH injections on Days 0, 3, 6 and 9. Subsequently, 78% of these synchronized animals responded to the PMSG stimulation and ovulated, with the mean number of collected oocytes increasing from 6 (control) to 20. This finding is particularly valuable when only limited numbers of female dunnarts are available for experiments.

In this study, we established a method and criteria to identity estrous stages in the dunnart using vaginal lavage cytology. Cyclic changes in epithelial cell morphology and leukocyte appearance within the vaginal and uterine lumen have been characterized in guinea pig, rat and mouse [17, 36-38]. The changes in the numbers and relative ratios of NECs, CECs and leukocytes reflect the fluctuations in circulating levels of the ovarian steroids (17-β-estradiol and progesterone) and gonadotropins (luteinizing and follicle stimulating hormones) throughout reproductive cycles. we employed a simple, non-invasive procedure to collect cell samples from the vaginal opening to avoid eliciting inflammatory responses or altering the reproductive cycle status. As demonstrated in this study, this technique enables the collection of sufficient cellular materials for precise determination of etrous cyclicity, with observable changes in cell morphology and composition directly reflecting underlying hormonal fluctuations and therefore the reproductive stages. This approach could be invaluable for broader application across marsupial species in managed captive settings, allowing to monitor breeding readiness, optimize mating timing, and enhance reproductive success by using assisted reproductive technologies for species preservation efforts, especially for threatened or endangered marsupial populations.

Finally, we also examined the developmental competence of oocytes derived from estrus synchronization and superovulation treatment. Zygotes were successfully collected from the adult females that underwent LHRH-PMSG treatments and mated with stud male dunnarts. These zygote embryos were collected approximately 5-6 hours post-hCG administration and developed to the blastocyst stage in dKSOM1/dKSOM2 sequential medium. These data indicate that oocytes generated through LHRH-PMSG combined treatment in adult female dunnarts have the potential to support early embryo development through to the blastocyst stage.

In conclusion, we have successfully established a high-yield superovulation protocol to generate mature MII oocytes by combining estrous cycle synchronization and PMSG ovarian follicle stimulation in adult female dunnarts. This protocol enables the effective use of adult female dunnarts and increases the yield of viable oocytes per animals that can be used in the development of ART and other experiments requiring large numbers of oocytes or embryos. This concept of combining estrus synchronization and follicle stimulation to yield sufficient MII oocytes or embryos may be applicable to a broad range of marsupial species to support captive breeding programs and ART assisted conservation efforts.

## Acknowledgements

The authors thank the staff at BioSciences 4 Animal Facility, the University of Melbourne, for the daily management of the dunnart colony. We acknowledge the Biological Optical Microscopy Platform at the University of Melbourne for the confocal microscopy. We acknowledge the Melbourne Histology Platform and Phenomics Australia Histopathology and Slide Scanning Service at the University of Melbourne for the assistance with tissue processing and imaging.

## Author contributions

JL̲design of the project, data collection and analysis, and manuscript writing and editing; NM̲data collection and analysis, and manuscript editing; ELS̲data analysis and manuscript editing; SO̲design of the project and manuscript editing; AJP̲design of the project, data analysis, and manuscript writing and editing.

## Data availability

Data are available on request.

